# Adult Clock Neuron Somatic Neuropeptide Release and Cytonemes Regulate Sleep

**DOI:** 10.64898/2026.07.27.741016

**Authors:** Markus K. Klose, Patricia K. Rivlin, Dinara Bulgari, Junghun Kim, Sydney N. Gregg, Xiju Xia, Yulong Li, Brigitte F. Schmidt, David L. Deitcher, Edwin S. Levitan

**Affiliations:** Department of Pharmacology and Chemical Biology, University of Pittsburgh, Pittsburgh, PA 15261, USA; Research and Exploratory Development, Johns Hopkins Applied Physics Laboratory, Laurel, MD 20723, USA; State Key Laboratory of Membrane Biology, School of Life Sciences, Peking University, Beijing 100871, China; PKU-IDG/McGovern Institute for Brain Research, Beijing 100871, China; Department of Chemistry, Carnegie Mellon University, Pittsburgh, PA, 15213, USA; Department of Neurobiology and Behavior, Cornell University, Ithaca, NY 14853, USA

## Abstract

*Drosophila* s-LNv clock neurons promote nighttime sleep by releasing the neuropeptide sNPF to activate sNPF receptors (sNPF-Rs) on l-LNv clock neurons. Behavior is controlled by synaptic transmission, but s-LNv and l-LNv neurons are not connected directly by chemical synapses. To investigate the basis of sNPF/sNPF-R communication between LNv neurons, the spread of sNPF was imaged in the adult brain. We report the daily midmorning burst of sNPF released from s-LNv terminals does not reach l-LNv neurons or s-LNv somata. Instead, sNPF released by the s-LNv soma late at night in response to sleep-promoting IP_3_ signaling reaches l-LNv somata, but not their terminals. In addition to communication by neuropeptide diffusion, analysis of fly connectomes revealed that adult s-LNv and l-LNv neurons form non-synaptic direct contacts mediated by cytonemes. Remarkably, genetically perturbing LNv neuron cytonemes alters sleep latency, but not nighttime sleep, the target of somatic sNPF release, or circadian behavior, which depends on PDF neuropeptide released by LNv terminals. Therefore, three distinct aspects of adult rhythmic behavior are produced by terminals, the soma and cytonemes, with the latter possibly acting via contacts that are not currently annotated in the connectome.

---

Imaging of a GPCR activation-based sensor for sNPF (encoded by UAS-GRAB_sNPF1.0_) (Xia and Li, 2025) expressed based on the PDF-GAL4 driver in brain explants (i.e., from PDF>GRAB flies) demonstrated s-LNv terminals release sNPF under control of the molecular clock at midmorning (i.e., ZT 3) (Klose *et al*., 2026). Explant confocal microscopy showed this sensor is also functional at LNv somata and l-LNv terminals (Fig. S1). However, l-LNv terminals failed to register any endogenous changes in sNPF at ZT 3 (the peak of release by s-LNv terminals) or other tested times (Fig. 1A, B). These results show that, even in the context of an insect’s small brain, the efficacious spread of neuropeptide released by terminals is limited.

**Figure 1.**
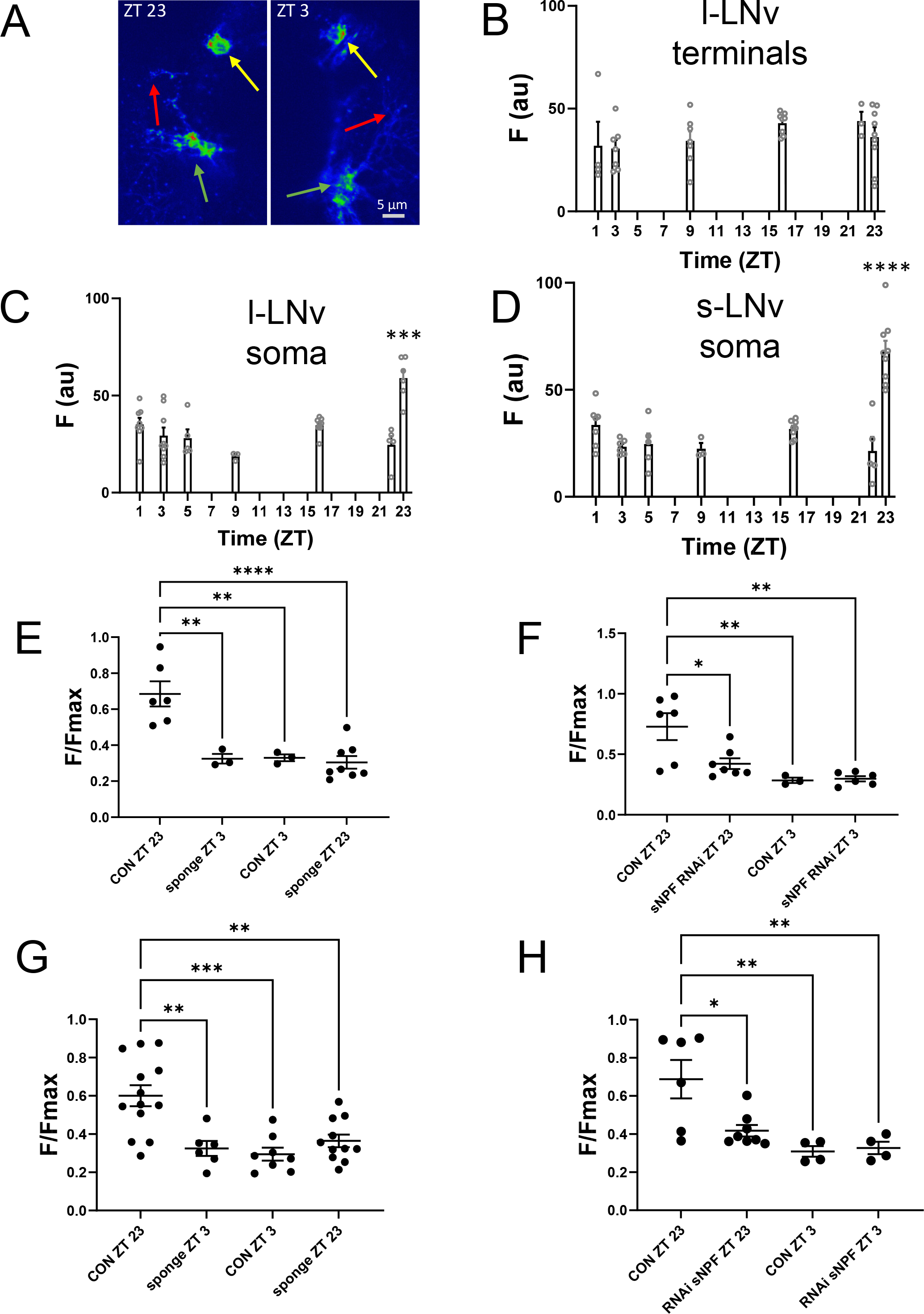
sNPF sensor signals in LNv neurons across the day of PDF>GRAB flies. A. sNPF sensor images in LNv neurons at ZT 23 and ZT 3. Bar, 5 µm. Orange arrows lLNv somata, green arrows sLNv somata, red arrow lLNv terminals. B. sNPF sensor fluorescence in l-LNv terminals does not change across the day. C. sNPF sensor fluorescence peaks daily in l-LNv somata at ZT 23. All points were different from data at ZT 23. Dunnett’s multiple comparison test. ***p < 0.001. D. sNPF sensor fluorescence peaks daily in s-LNv somata at ZT 23. All points were different from data at ZT 23. Dunnett’s multiple comparison test. ****p < 0.0001. E. The daily peak in sNPF sensor fluorescence in s-LNv somata was absent in UAS-IP_3_ sponge/ PDF>GRAB flies. F. The daily peak in sNPF sensor fluorescence in s-LNv somata was absent in UAS-sNPF RNAi/PDF>GRAB flies. G. The daily peak in sNPF sensor fluorescence in l-LNv somata was absent in UAS-IP_3_ sponge/PDF>GRAB flies. H. The daily peak in sNPF sensor fluorescence in l-LNv somata was absent in UAS-sNPF RNAi/PDF>GRAB flies. For E-H, Tukey’s multiple comparisons tests were used. **p < 0.01, ***p < 0.001, ****p < 0.0001.

We then examined endogenous sNPF at l-LNv somata, which are responsive to sNPF (Shang *et al*., 2013), but do not contain sNPF (Johard *et al*., 2009). First, no sNPF was detected at l-LNv somata at ZT 3, but a response was detected at ZT 23 (Fig. 1A, C). Prior experiments showed neuropeptide-containing dense-core vesicles (DCVs) in the s-LNv soma undergo exocytosis at ZT 23 in response to IP_3_-IP3R signaling that increases sleep consolidation (Klose *et al*., 2021). Therefore, we quantified sNPF sensor signals to test if daily DCV exocytosis at s-LNv somata releases sNPF that could support the l-LNv response. These experiments demonstrated native extracellular sNPF levels at s-LNv somata do not change at ZT 3, further demonstrating neuropeptide release from terminals does not reach LNv somata. However, there is an sNPF increase at s-LNv somata at ZT 23 (Fig. 1A, D). Furthermore, this effect is inhibited by cell-specific UAS-IP_3_ sponge expression or UAS-RNAi-mediated knockdown of sNPF (Fig. 1E, F). Thus, native sNPF is released from s-LNv somata at ZT 23 based on IP_3_ signaling. Finally, these genetic perturbations also blocked the sNPF signal at l-LNv somata at ZT 23 (Fig. 1G, H). Therefore, sNPF released by s-LNv somata selectively reaches l-LNv neurons at their somata. Given that IP_3_ signaling promotes sleep and triggers sNPF release by s-LNv somata (Klose *et al*., 2021, Fig. 1E) and l-LNv somata alter their intracellular signaling in response to sNPF (Shang *et al*., 2013), the above results show that the sleep controlling sNPF-based communication between s-LNv and l-LNv neurons is based on soma-to-soma peptidergic transmission.

*Drosophila* brain neurons are typically unipolar with a proximal neurite extending from the soma that gives rise to the dendritic arbor and the axon. However, a subset of CRY-expressing evening clock neuron somata in fixed preparations also feature membrane outgrowths (Schubert *et al*., 2018). Because such structures could participate in communication between clock neurons, we considered whether outgrowths protrude from s-LNv and l-LNv somata. Reconstructions derived from multiple *Drosophila* connectomic datasets (Scheffer *et al.,* 2020; Nern *et al.,* 2025; Berg *et al.,* 2025) show s-LNv and l-LNv somata have outgrowths that are thin compared to their proximal neurites (Fig. 2A). For example, the longest outgrowth in the shown s-LNv neuron (Fig. 2A, left) is ∼20 µm in length with a diameter that varies from 150-400 nm, which is reminiscent of cytonemes in the imaginal disc (Ramirez-Weber and Kornberg 1999). These outgrowths were also apparent with live optical imaging in the adult brain explant (Fig. 2B-G). In some cases, they were highly dynamic (Fig. 2B, Movies S1 and S2), which is again reminiscent of cytonemes (Ramirez-Weber and Kornberg 1999; Daly *et al*., 2022). Furthermore, indicative of a signaling function, cell specific expression of Dilp2-GFP, a dense-core vesicle (DCV) marker (Wong *et al*., 2012), and Dilp2-FAP, an indicator of DCV kiss and run exocytosis (Bulgari *et al*., 2019; Bulgari *et al*., 2023), revealed DCVs are present in living s-LNv outgrowths and undergo exocytosis (Fig. 2D,E). Outgrowths are also labeled by moesin-GFP, a marker for actin (Fig. 2F). Taken together, the above features are suggestive of signaling filopodia (i.e., cytonemes), which have been studied in *Drosophila* during development (Ramirez-Weber and Kornberg 1999; Kornberg and Roy, 2014; Daly *et al.,* 2022). Finally, cell specific expression of *diaphanous* (*dia*) RNAi, which shortens *Drosophila* cytonemes (Roy *et al*., 2014), reduced the length of the longest outgrowth from s-LNv somata (Fig. 2G,H). Therefore, the above data show adult LNv clock neuron somata possess membranous outgrowths called cytonemes.

**Figure 2.**
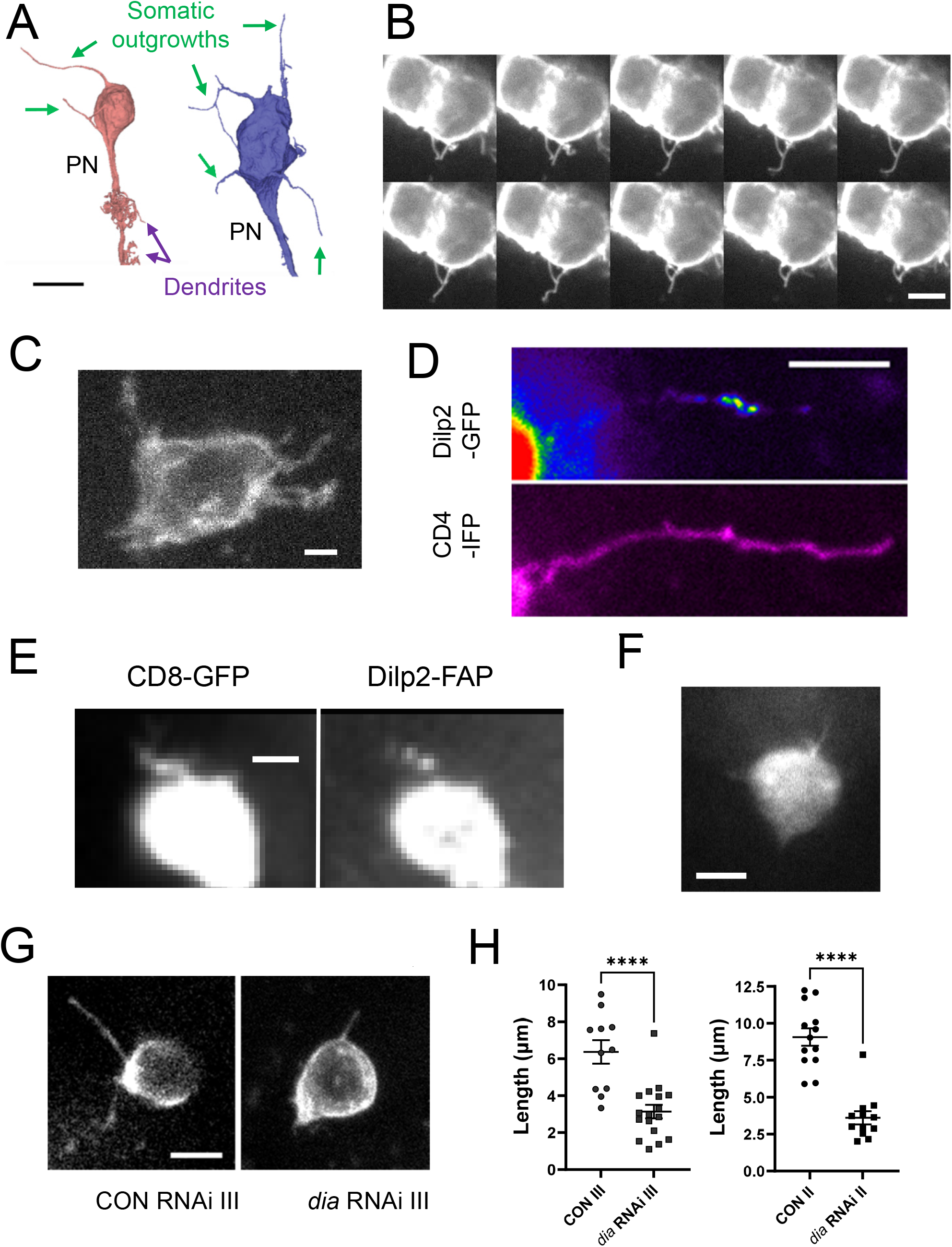
Cytonemes emanate from LNv somata. A. 3D reconstructed s-LNv (red) and l-LNv (blue) somata from connectome datasets. The proximal neurite (PN) in each case was oriented down. Size bar, 10 µm. B. Images from a live brain explant show l-LNv somata in PDF>GRAB animals can possess thin and dynamic outgrowths. Images were acquired at 0, 10, 170, 180, 190, 220, 230, 340, 350, and 360 seconds of a time-lapse experiment. Size bar, 5 µm. For complete time-lapse experiment, see Movie S1. C. Brain explant image of s-LNv soma. Size bar, 2 µm. D. CD4-mIFP reveals outgrowth from an s-LNv soma with DCVs labeled with Dilp2-GFP. Size bar is 5 µm. E. Co-expression of CD8-GFP and Dilp2-FAP reveals DCV exocytosis in an s-LNv outgrowth. Size bar, 2 µm. F. s-LNv outgrowths in PDF>GRAB animals are labeled with moesin-GFP. Size bar, 4 µm. See Movie S2 shows dynamics. G. s-LNv outgrowths in in PDF>GRAB animals with a control RNAi (CON RNAi III) and with *diaphanous* RNAi expression (*dia* RNAi III, see Methods). Size bar, 5 µm. H. Quantification of the length of the longest s-LNv cytonemes with two *dia* RNAis (*dia* RNAi III and RNAi II) and their matched controls (CON RNAi III and II, see Methods). ****p < 0.0001, unpaired t-test.

Prior connectome analyses showed that s-LNv and l-LNv neurons are not connected by direct conventional synapses (Shafer *et al*., 2022, Reinhard *et al*., 2024), but the possibility that cytonemes could form contacts between these clock neurons was not considered. To test for contacts between LNv neurons, we initially conducted trans-Tango experiments (Talay *et al*., 2017). With an updated version of this assay (see Methods), either s-LNv or l-LNv cells expressed CD8-GFP and a tethered ligand based on a split-GAL4 driver, while mtd-Tomato was expressed with trans-Tango in contacting neurons. To further identify s-LNv and l-LNv neurons, the neuropeptide PDF was immunolabeled with a far-red fluorophore-labeled secondary antibody. Such triple labeling showed s-LNv neurons form contacts amongst themselves and with l-LNv neurons (Fig. 3Ai). In complementary experiments, l-LNv neurons contacted themselves and s-LNv neurons (Fig. 3Aii). Such extensive contacts between s-LNv and l-LNv neurons were seen in all brain hemispheres (N = 12) that were examined. Finally, ultrastructural features resolved in the connectomic EM volumes were used to determine the structural basis of these contacts (Fig. 3B). These studies verified that s-LNv cytonemes contain DCVs that undergo exocytosis, as is evident from omega profiles. Furthermore, in contrast to dendrites and axons, microtubules were not observed in cytonemes. In addition, s-LNv and l-LNv cytonemes were found to form direct membrane-to-membrane contacts that are ensheathed by glia (Fig. 3B, right panels). Importantly, these contacts were devoid of conventional synaptic markers (e.g., T-bars, post-synaptic densities, small clear vesicle clusters). Together, the above results demonstrate s-LNv and l-LNv neurons are connected by non-synaptic peptidergic cytoneme-based contacts that are not annotated by conventional connectome analysis.

**Figure 3.**
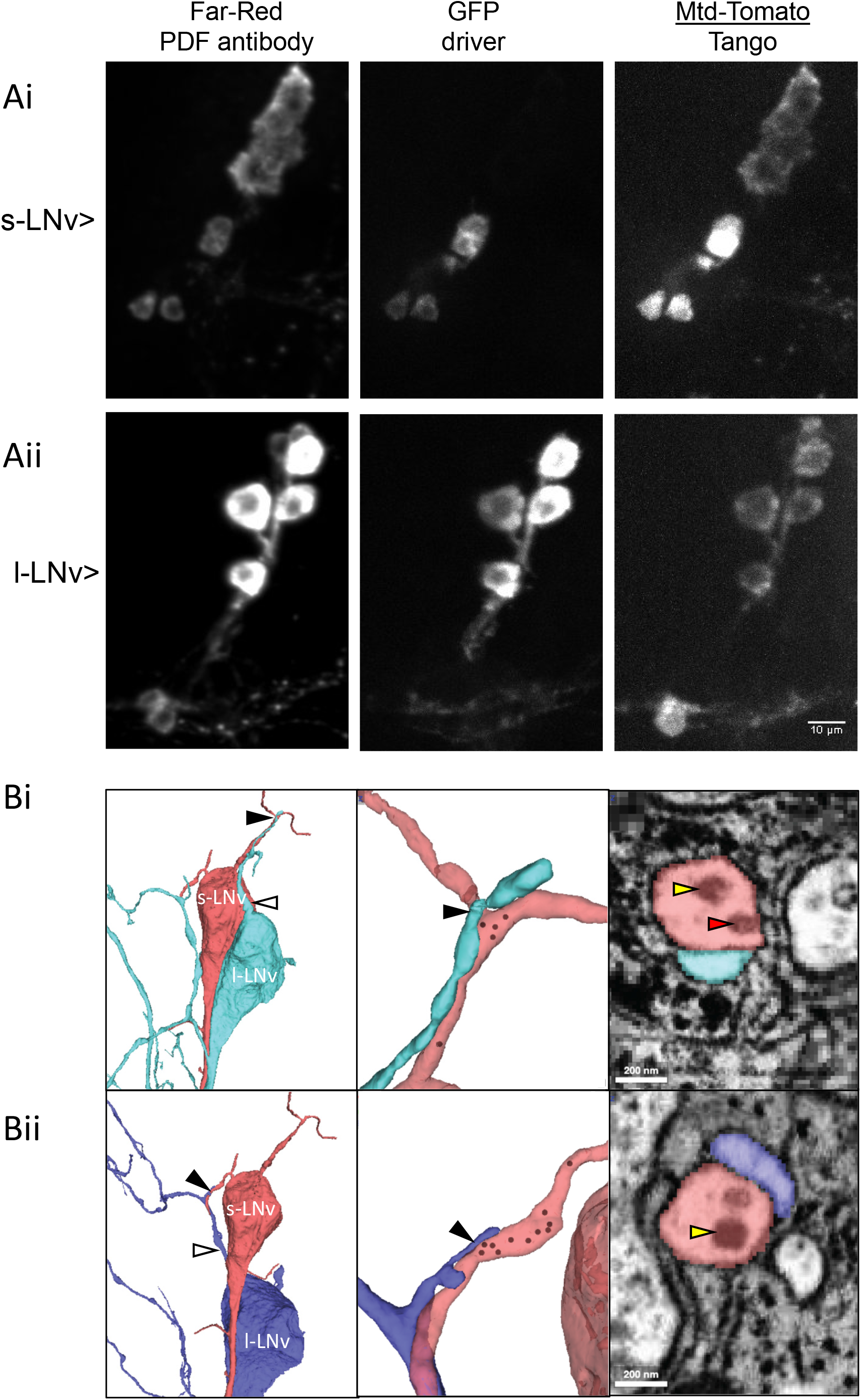
Trans-Tango and ultrastructure show contacts are made between LNv neurons. Ai, left panel shows both s-LNv and l-LNv neurons labeled for PDF content; middle, s-LNv split-GAL4 expression produces GFP expression in s-LNv neurons; right, Tango expression, which indicates contacts, is in s-LNv neurons and l-LNv neurons. Aii, l-LNv split-GAL4 expression of TANGO produces GFP expression in l-LNv neurons and Tango expression in both s-LNv and l-LNv neurons. Size bar, 10 µm. B. 3D reconstructions of a single s-LNv (red), and two l-LNv neurons (light blue and dark blue). Left panels: l-LNv cytonemes (white arrows) appear to make direct contacts with s-LNv cytonemes (black arrows). Center panels: zoom-in view of s-LNv-l-LNv contact site (black arrow) reveals DCVs (annotated with black dots in 3D reconstruction) within s-LNv cytonemes. Right panels: Electron microscopy images of cytonemes in cross section reveal direct sLNv–lLNv contacts, DCVs (yellow arrows), and a putative omega profile (red arrow). Note that cytonemes are ensheathed by glia (dark cytoplasm) with no glial processes detected between them. Bars, 200 nm.

We next addressed whether s-LNv and l-LNv cytonemes influence behaviors associated with these clock neurons. Specifically, *dia* RNAi was expressed in LNv neurons under the control of PDF-GAL4 and locomotor activity was monitored under both LD illumination (12 hour light:12 hour dark) and then in constant darkness (DD) to test for effects on sleep and clock function. No changes in circadian locomotor behavior were evident: activity, period and rhythmicity did not change and morning anticipation was still evident (Fig. 4A-C). Thus, *dia* knockdown does not affect synaptic control of the clock circuit by LNv neurons, which relies on activity-dependent release of the neuropeptide PDF from terminals (King and Sehgal, 2018).

**Figure 4.**
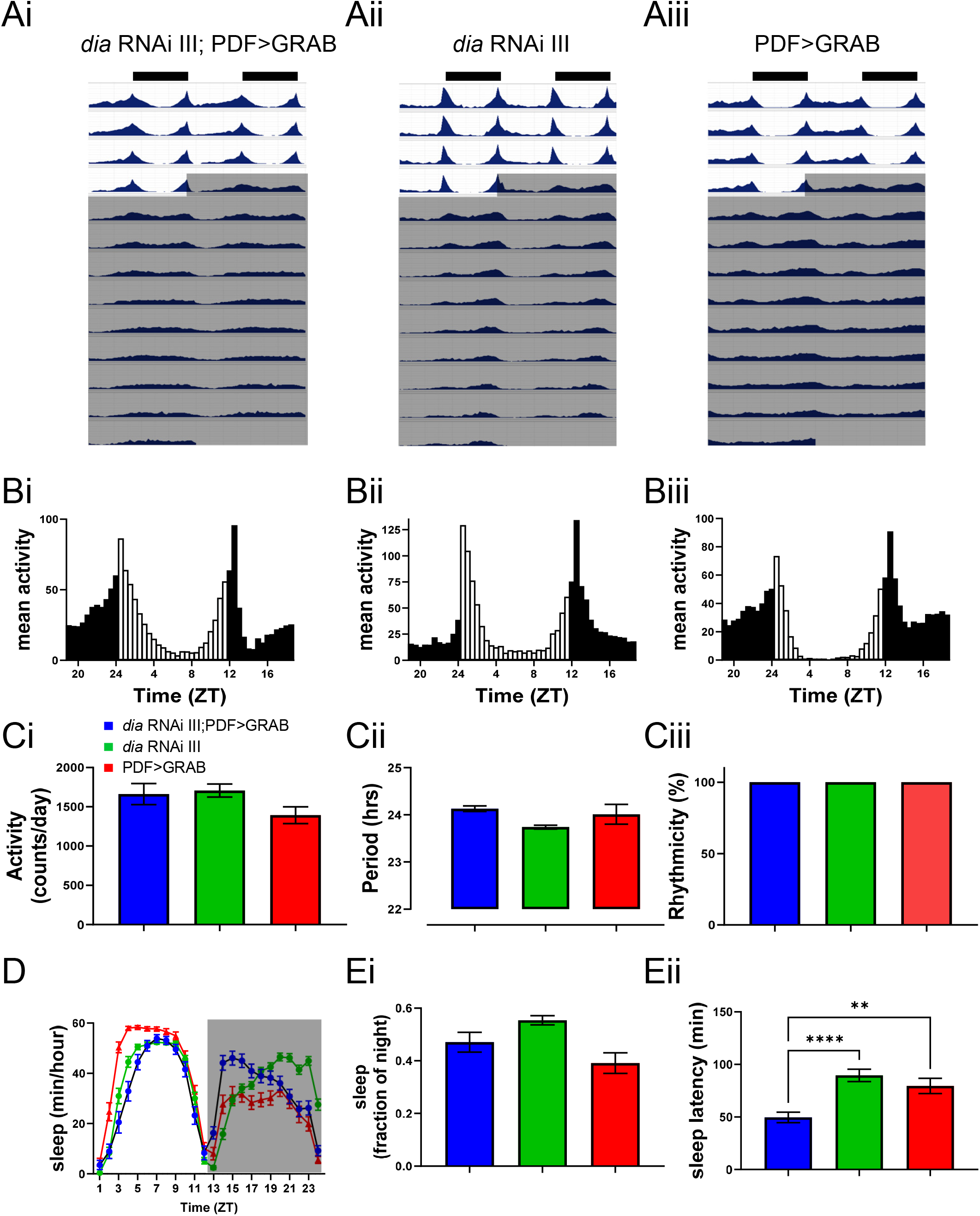
*Diaphanous* RNAi (*dia* RNAi) expressed in LNv neurons reduces sleep latency without affecting total sleep or the circadian rhythm. A. Actograms of flies expressing *diaphanous* RNAi in PDF neurons (*dia* RNAi III; PDF>GRAB, left) compared to two genetic controls, (*dia* RNAi without a driver, middle and PDF>GRAB, right) Flies entrained under a 12 h:12 h light/dark regimen (LD) before being shifted to DD. Data shown are from one experiment with 32 flies in each genotype. B. Eductions of activity across the day in LD cycle. Ci. Activity counts per day (average of last two days in LD). Cii. Period of daily behavioral rhythms in DD cycle (calculated using day 2-9 of DD). Ciii. Percent of flies that are rhythmic (calculated using day 2-9 of DD). D. Sleep in flies expressing *diaphanous* RNAi in PDF neurons (*dia* RNAi III; PDF>GRAB) compared to two genetic controls, *dia* RNAi1 and PDF>GRAB). Sleep presented as minutes per hour across the day. Data shown are from one experiment with 32 flies in each genotype (average of the last two days in LD). Ei. Expression of *diaphanous* RNAi1 in PDF neurons had no effect on total sleep. Eii. Sleep latency in flies expressing *diaphanous* RNAi1 in PDF neurons (Dia RNAi1; PDF>GRAB) compared to two genetic controls, *dia* RNAi1 and PDF>GRAB. All tests were One-Way ANOVAs with Dunnett’s multiple comparison test. **p < 0.01, ****p < 0.0001. See Fig. S2 for related data.

Next we analyzed the effect of *dia* knockdown on sleep, which is promoted by sNPF released by s-LNv neurons (Shang *et al*., 2013) and the IP_3_ signaling that evokes somatic sNPF release by s-LNv somata and promotes sleep (Klose *et al*., 2021; Fig. 1). No changes occurred in nighttime sleep, but sleep latency was reduced (Fig. 4D,E) regardless of expression of the GRAB sensor, the use of two different *dia* RNAis and RNAi insertion site controls (Fig. S2). Because past studies did not examine the effect of LNv sNPF/sNPFR signaling on sleep latency, sleep behavior was quantified after cell specific Crispr-Cas9 knockout the sNPF-R gene. These experiments verified the previously reported effect of sNPF/sNPF-R signaling on nighttime sleep, but no effect on sleep latency was observed (Fig. S3). Thus, the behavioral effect of LNv cytonemes differs from the known effect of sNPF, thereby demonstrating an sNPF-independent action of cytonemes.

Connectomics is pursued with the goal of explaining behavior in terms of neural circuits and their synaptic connectivity in the brain. However, clock neuron studies illustrate limitations of focusing exclusively on conventional chemical synapses. First, somata of two different clock neurons (s-LNv and l-LNv) communicate by neuropeptide diffusion instead of direct synapses to promote nighttime sleep. Second, LNv clock neuron cytonemes influence a specific aspect of sleep (latency) that is unaffected by sNPF, consistent with control of adult behavior by cytoneme contacts. This effect is also distinct from the control of circadian rhythms by PDF released from LNv terminals (King and Sehgal, 2018). Therefore, each of three different compartments (terminals, somata and cytonemes) of adult clock neurons regulates its own unique behavioral feature. This remarkable conclusion indicates that explaining the neural basis of behavior will benefit by incorporating cytonemes and their contacts, as well as the proximity of peptidergic neuronal somata, into connectome analysis.

## Methods

### Optical Imaging

Dissections were performed as previously described with anterior of brain explants facing upwards (Klose *et al*., 2026). Adult flies were immobilized with CO_2_ gas and brains were dissected in 0 Ca^2+^ HL3 saline solution (70 mM NaCl, 5 mM KCl, 20 mM MgCl_2_•6 H_2_O, 115 mM Sucrose, 5 mM Trehalose, 5 mM Hepes, and 10 mM NaHCO_3_, pH 7.3) and then put into polylysine-coated plastic dishes containing HL3 with 2 mM Ca^2+^ for imaging. All imaging was done on setups with upright Olympus microscopes equipped with dipping water immersion objectives (60 or 20×), Yokogawa spinning disk confocal heads, lasers (488 nm laser for GFP, 561 nm laser for RFPs and a 640 nm laser for imaging of far-red fluorescence in FAP and immunofluorescence) and a Teledyne Photometrics sCMOS camera. Fluorescent probes included moesin-GFP, CD8-GFP and CD4-mIFP membrane markers, mtd-Tomato, the sNPF sensor GRAB_sNPF1.0_ and its mutant (Xia and Li, 2025), and a far-red-labeled secondary antibody (Alex Fluor 647 Conjugated affinipure Goat Anti-mouse IgG (H+L), Jackson ImmunoResearch), which was used with a primary monoclonal antibody against PDF (PDF C7, Developmental Studies Hybridoma Bank). As in our previous study (Klose *et al*., 2026), UAS-GRAB_sNPF1.0_ expression in LNv neurons was driven by PDF-GAL4, here denoted as PDF>GRAB, or split-GAL4 drivers (see below). Sensor function was demonstrated by application of sNPF2 (WFGDVNQKPIRSPSLRLRFamide) (GenScript Life Science). FAP imaging experiments were performed as previously described (Klose et al., 2021). Fluorescence was quantified in ImageJ or Fiji. Analysis of cytoneme length was limited to examples in which cytonemes were well resolved in a single plane of focus. Many other images are projections from multiple planes of focus.

### Fly lines

Previously reported flies included UAS-Dilp2-GFP/CyO (Wong *et al*., 2012), UAS-Dilp2-FAP/CyO (Bulgari *et al*. 2019) and UAS-GRAB_sNPF1.0_ and its mutant (Xia and Li, 2025). The PDF-GAL4 driver on the third chromosome (provided by Paul Taghert, Washington University in St. Louis) was used to drive expression in the two subsets of clock neuron that express the PDF neuropeptide, the small ventrolateral (s-LNv) neurons and the large ventrolateral (l-LNv) neurons, of which only the s-LNv neurons express sNPF. C. Andrew Frank (University of Iowa) and Leslie Griffith (Brandeis University) provided fly lines with chromosome 3 insertions of UAS-IP_3_ sponge and UAS-sNPF RNAi flies, respectively. For sNPF-R cell specific knockout, a gRNAx3 construct line (P{UAS-sNPF-R.gRNAx3.pCFD6}attP1) (Schlichting *et al*., 2022, Proc. Natl. Acad. Sci. U.S.A. 119(34): e2206066119) was provided by Michael Rosbash (Brandeis University). Trans-Tango experiments used w; trans-Tango/CyO; QUAS-mtdTomato, UAS-CD8-GFP/TM6,Tb flies that were generated by crossing a recombinant of Bloomington *Drosophila* stock center fly lines BDSC# 30005 and 32187 on chromosome III to the trans-Tango insertion on chromosome II (BDSC# 77123).

Other BDSC lines used here include s-LNv split-GAL4 (BDSC# 87860), l-LNv split-GAL4 (BDSC# 87861), UAS-moesin-GFP (P{w[+mC]=UAS-GMA}3 (BDSC# 31776), P{y[+t7.7] w[+mC]=UAS-CD4-mIFP-T2A-HO1}attP40 (BDSC# 64183), two *dia* UAS-RNAi lines that are referred to (based on their chromosome insertion sites) as RNAi III (BDSC# 33424) and RNAi II (BDSC #80437) and their respective chromosome insertion site controls (BDSC# 35785, CON III and 67852, CON II). Finally, BDSC# 67077 was used to generate UAS-Cas9.P2; PDF-GAL4.

### Quantification and statistical analysis

Statistical analyses were performed using Prism (GraphPad). Bar graphs show means with standard error of the mean indicated by error bars, as well as individual data points. Comparisons of multiple experimental groups were based on one-way ANOVA followed by Tukey’s or Dunnett’s multiple comparisons post-tests. Pair-wise analysis used unpaired t-tests. Statistical significance is indicated when two-tail p values were ≤ 0.05.

### Ultrastructural Analysis of Cytonemes

We manually examined s- and l-LNv neurons for the presence of cytonemes in each brain hemisphere of the Hemibrain, Optic Lobe, and Male Central Nervous System (MCNS) connectomic datasets (Scheffer *et al.,* 2020; Nern *et al.,* 2025; Berg *et al.,* 2025). Briefly, the Hemibrain dataset contains the central brain from a 5-day-old adult female fly, the MCNS dataset contains the entire brain and ventral nerve cord from a 5-day-old adult male fly, and the Optic Lobe dataset comprises the right optic lobe reconstructed from the MCNS sample. All datasets were acquired using Focused Ion Beam Scanning Electron Microscopy (FIB-SEM) and reconstructed using automated segmentation followed by manual proofreading to correct segmentation errors.

Although cytonemes have been reported to be poorly preserved following conventional chemical fixation (Ramirez-Weber and Kornberg, 1999), they are well preserved in these FIB-SEM datasets, which were generated using tissue preparation protocols optimized for connectomics (Lu *et al.,* 2022). Together with the isotropic voxel resolution of FIB-SEM, these preservation methods make the Hemibrain, Optic Lobe and MCNS datasets well suited for ultrastructural analysis and tracing of cytonemes.

For each dataset, the EM image volume and neuron segmentations were visualized in 3D using *Neuroglancer*, available through the *NeuPrint Connectome Explorer* (Plaza *et al.,* 2022) at neuprint.janelia.org. Because cytonemes are thin processes (∼200 nm in diameter in the imaginal disc; Wood *et al*., 2021) and can extend for tens of micrometers, many cytonemes are only partially reconstructed in the publicly available Hemibrain and MCNS segmentations. Therefore, for each LNv soma, we identified the distal tips of cytoneme-like structures and short membrane protrusions that could represent incompletely reconstructed cytonemes. We then systematically proofread each candidate structure, extending it to its distal endpoint whenever supported by the underlying EM data.

### Behavior

Entrained adult flies (3 to 4 d old) were loaded into glass tubes and placed in DAM2 Trikinetics Activity Monitors. Flies were maintained on a 12 h:12 h light:dark (LD) schedule for 7 days and then released into constant darkness for 9 days. We assessed rhythmicity by normalizing activity from the 2^nd^ to 9^th^ day of constant darkness (12 h: 12 h dark:dark). We defined arrhythmic flies by rhythmicity threshold [Qp.act/Qp.sig] below 1 or a period estimate <18 or >30 h. Activity, rhythmicity and period were assessed using ShinyR-DAM software (Cichewicz and Hirsh, 2018). Sleep parameters were derived using a custom analysis program as used in Shaw *et al*. (2000).

## Acknowledgements

We thank Drs. C.A. Frank, Leslie Griffith and Michael Rosbash for providing flies. Stocks obtained from the Bloomington Drosophila Stock Center (NIH P40OD018537) were used in this study. The PDF C7 monoclonal antibody developed by Justin Blau was obtained from the Developmental Studies Hybridoma Bank, created by the NICHD of the NIH and maintained at The University of Iowa, Department of Biology, Iowa City, IA 52242. We also thank the FlyEM Project at the Janelia Research Campus for access to the Optic Lobe and MCNS datasets and Stuart Berg for assistance with proofreading. Research reported in this publication was supported by the National Institute of Neurological Disorders And Stroke of the National Institutes of Health under Award number R01NS032385 to ESL. The content is solely the responsibility of the authors and does not necessarily represent the official views of the National Institutes of Health. PR was additionally supported by the National Institute of Mental Health of the National Institutes of Health under Award Number R24MH114785.

---

**Figure S1.**
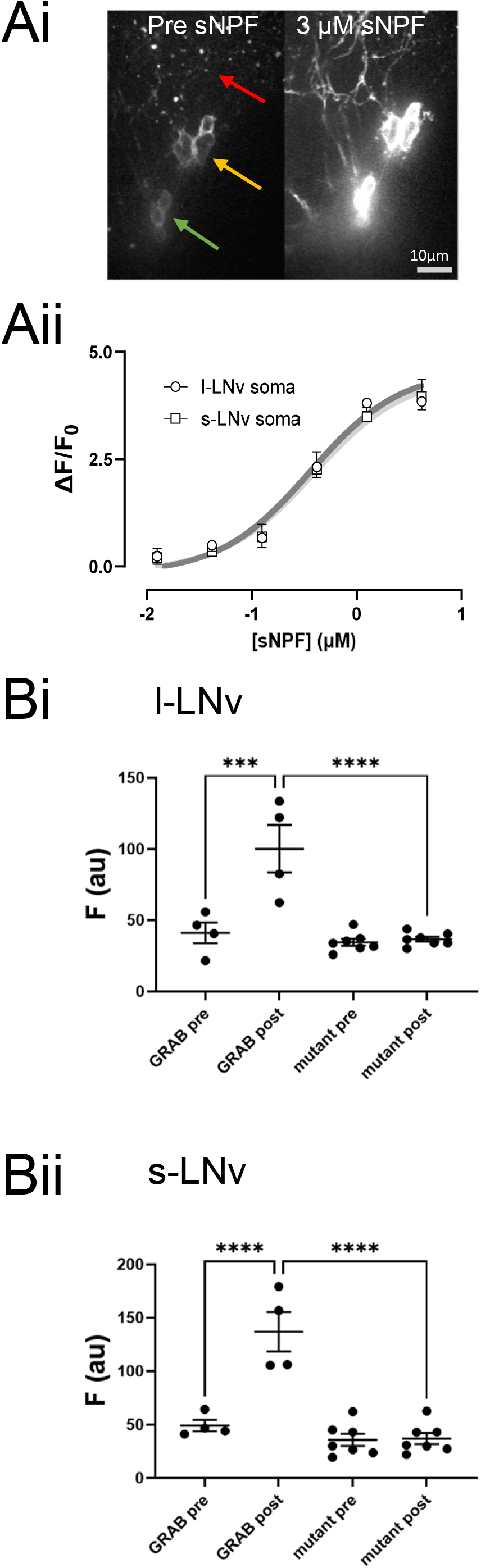
GRABsNPF1.0 in LNv somata responds to exogenous sNPF2 neuropeptide in a dose-dependent manner. Ai. Images of GRAB_sNPF1.0_ fluorescence in LNv neurons before (Pre) and after exogenous application of 3 µM sNPF2 (red arrow-l-LNv terminals, yellow arrow-l-LNv somata, green arrow-s-LNv somata). Images of ventro-lateral brain and optic lobe in PDF>GRAB adult flies before and after 3 µM exogenous sNPF2 was applied. Regions of interest were drawn around all somata and measured for GRAB sensor intensity. Aii. Dose-response curves of the GRAB sensor in l-LNv somata and s-LNv somata. Bi. Unlike the normal sensor (GRAB pre and post), a nonfunctional mutant sensor mutant (mutant) in l-LNv somata does not respond to exogenous application of 3 µM sNPF2 (mutant post). Bii. A nonfunctional sensor mutant also does not respond to exogenous sNPF2 application in s-LNv somata.

**Figure S2.**
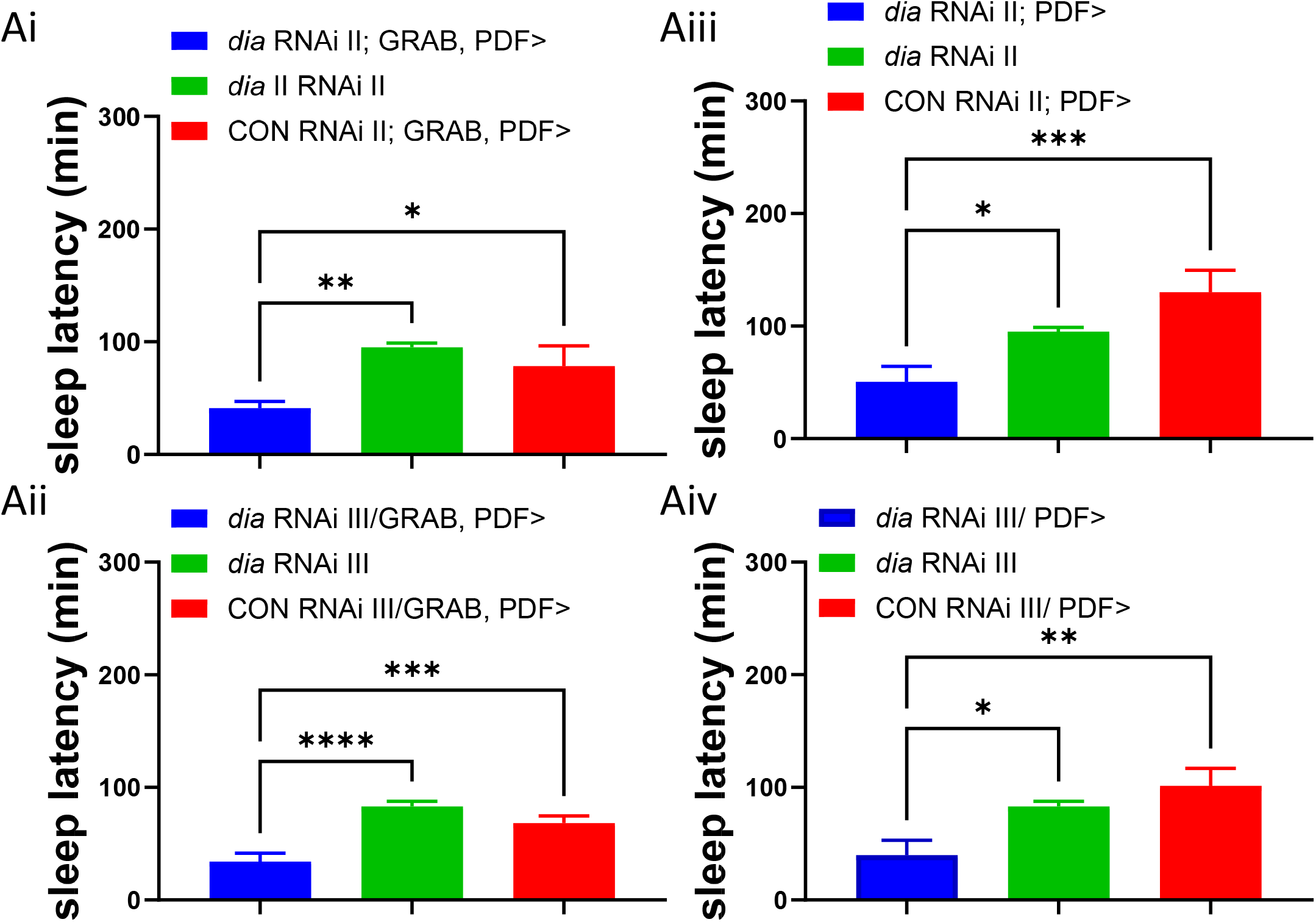
Effects on sleep latency of two *dia* RNAis on chromosomes II and III compared to insertion site RNAi controls (CON RNAi II and III, see Methods) with the GRAB sensor (Ai and Aii) and without the GRAB sensor (Aiii and Aiv). Data were analyzed by one-way ANOVA and Dunnett’s multiple comparison test., *p < 0.05, **p < 0.01, ****p < 0.0001.

**Figure S3.**
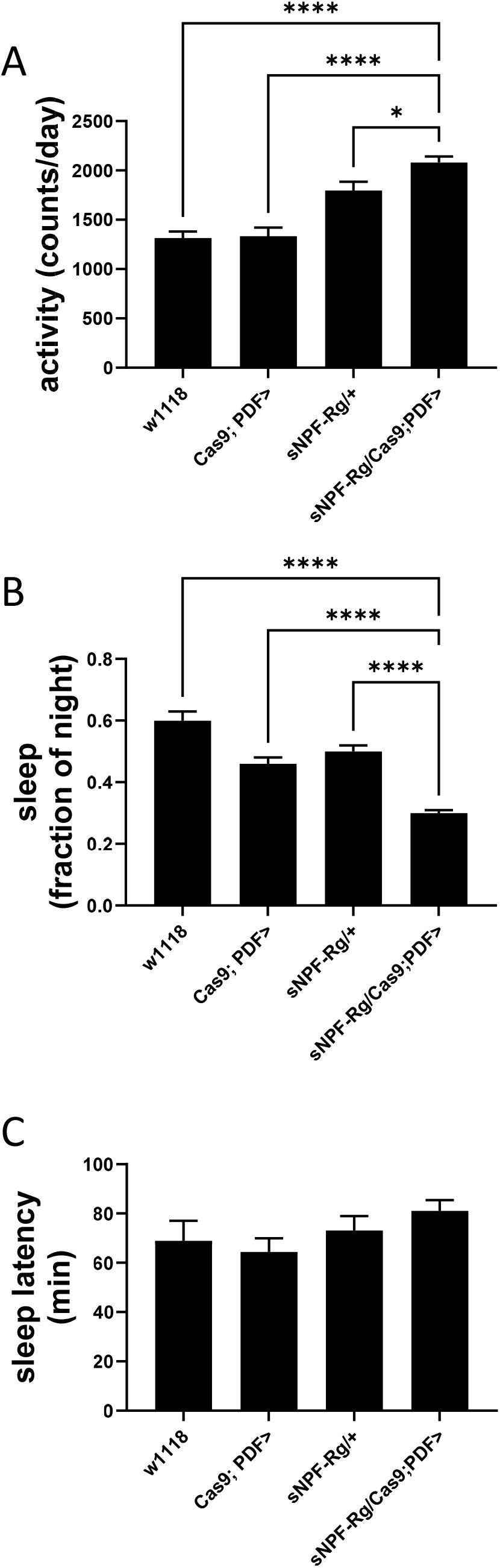
sNPF-R knockout in LNv neurons decreases nighttime sleep and daily locomotor activity without affecting sleep latency. A Activity in flies expressing sNPF-R gRNA (sNPF-Rg) in PDF neurons (sNPF-R g/Cas9; PDF>) compared to two genetic controls, sNPFRg/+ and Cas9; PDF>. Flies entrained under a 12 h:12 h light/dark regimen. Data shown are from one experiment with 32 flies in each genotype. B. Expression of sNPF-R gRNA in PDF neurons with Cas9 decreased nighttime sleep. C. Expression of sNPF-R gRNA in PDF neurons with Cas9 did not affect sleep latency. Dunnett’s multiple comparison tests were used. **p < 0.01, ****p < 0.0001.

**Movie S1. Dynamic cytonemes are shown from on l-LN somata of a PDF>GRAB fly.** The time-lapse movie was acquired at 0.1 Hz for 60 seconds.

**Movie S2. Time-lapse movie from s-LNv soma expressing moesin-GFP.** Images were acquired at 10 Hz for 15 seconds.

